# A Relaxin-Defined Late-Luteal Endocrine Transition State in the Human Menstrual Cycle

**DOI:** 10.64898/2026.10.25.754541

**Authors:** Yasin Dhaher, Subaryani Soedirdjo, Sungsoo Lee, Luis A. Rodriguez, Ellen Casey

**Affiliations:** Department of Physical Medicine and Rehabilitation, University of Texas Southwestern Medical Center, 5323 Harry Hines Blvd, Dallas, TX 75390, United States; Department of Bioengineering, University of Texas at Dallas, 800 W Campbell Rd, Richardson, TX 75080, United States; Peter O’Donnell Jr. Brain Institute, University of Texas Southwestern Medical Center, 5323 Harry Hines Blvd, Dallas, TX 75390, United States; Department of Physiatry, Hospital for Special Surgery, New York, New York, United States

## Abstract

Dynamic endocrine transitions govern female physiology, yet several reproductive hormones remain incompletely characterized across the natural menstrual cycle. Relaxin, a peptide hormone classically associated with pregnancy, has recognized vascular, extracellular matrix, and anti-fibrotic actions, but its physiology in nonpregnant women remains poorly defined. We performed dense longitudinal endocrine profiling across 80 natural menstrual cycles from 34 healthy women, complemented by oral contraceptive users, to define circulating relaxin-2 dynamics relative to ovarian steroids.

Using generalized additive modeling of cycle-normalized trajectories, we identified relaxin as a highly phase-locked late-luteal hormone with a sharp, reproducible peak distinct from estradiol and progesterone. Progesterone rose earlier after ovulation and peaked at cycle day 23.9, whereas relaxin peaked significantly later at day 28.0, revealing an approximately 4-day temporal offset. Relaxin showed lower inter-cycle variability and narrower temporal dispersion than progesterone, indicating unexpectedly precise endocrine timing. Detectable relaxin persisted into early menses, suggesting continued luteal peptide release across the menstrual boundary.

In combined oral contraceptive users, endogenous cyclicity was attenuated and circulating relaxin was effectively absent despite exogenous progestin exposure, implicating ovulation-dependent corpus luteum biology rather than steroid exposure alone as the primary determinant of relaxin secretion.

These findings identify relaxin as a reproducible component of normal cyclic endocrine physiology rather than a hormone restricted to pregnancy. The discrete late-luteal relaxin window establishes a previously under-resolved temporal feature of the menstrual cycle and provides a framework for investigating its reproductive, vascular, and connective-tissue significance.

## Introduction

Relaxin, a peptide hormone of the insulin-like family, is best known for its reproductive functions during pregnancy, where it promotes cervical ripening, pelvic remodeling, uterine quiescence, and maternal cardiovascular adaptations ^1,2^. The principal human isoform, relaxin-2, acts through the G protein–coupled receptor RXFP1, which is expressed in reproductive, vascular, renal, and musculoskeletal tissues ^3,4^. Beyond pregnancy, relaxin contributes to extracellular matrix turnover, vasodilation, and anti-fibrotic signaling ^5,6^, suggesting broader physiological roles that extend beyond gestation.

Despite extensive study of relaxin in pregnancy, its secretion dynamics and physiological function during the menstrual cycle remain poorly understood. Circulating relaxin is detectable at low concentrations in non-pregnant women, yet its temporal relationship with ovarian steroid hormones has not been systematically defined. Early studies report variable or inconsistent cyclic patterns^7–9^, which likely reflect differences in sampling resolution, assay sensitivity, and interindividual variation in corpus luteum function. A clearer understanding of these dynamics is essential, as the corpus luteum co-secretes both progesterone and relaxin, which may act in a coordinated sequence to regulate luteal function, vascular remodeling, and extracellular matrix dynamics.

Progesterone synthesis begins immediately after ovulation, stimulated by the luteinizing hormone (LH) surge, which activates cholesterol conversion pathways in luteal cells. This rapid steroidogenic response leads to an early rise in serum progesterone, initiating endometrial secretory transformation and supporting potential implantation. In contrast, relaxin synthesis involves gene transcription and translation of the RLN2 gene, followed by peptide processing and regulated secretion within the corpus luteum. Its release appears to depend on the degree of luteal vascularization and perfusion, which typically peak later in the mid-luteal phase ^1,2^. As a result, relaxin exhibits a delayed serum rise relative to progesterone. Functionally, this temporal offset may be adaptive: progesterone prepares the endometrium for metabolic and structural readiness, while relaxin fine-tunes extracellular matrix remodeling, angiogenesis, and vasodilation to sustain luteal stability and support implantation potential.

Understanding this relationship has implications beyond reproductive physiology. Relaxin’s effects on connective tissue turnover, ligament compliance, and vascular tone suggest possible links between menstrual cycle–dependent hormone fluctuations and musculoskeletal function. Elevated relaxin levels in the luteal phase have been associated with increased joint laxity and ligament injury risk, although findings are inconsistent and may depend on concurrent progesterone and estrogen levels ^10,11^. Establishing the precise temporal and quantitative profile of relaxin across the menstrual cycle, alongside progesterone, will thus clarify both its endocrine coordination and its potential contribution to cyclic tissue remodeling.

The present study aims to define the circulating profile of relaxin-2 across the menstrual cycle and to determine its temporal relationship with progesterone. By delineating their relative timing and amplitude, this work seeks to provide a physiological foundation for understanding relaxin’s role in luteal phase biology and its relevance to women’s health.

## Methods

Thirty-four healthy women (age: 27 ± 5 yr.; BMI: 25.4 ± 5.6 kg/m2; means ± SD), non-users of hormonal contraceptives, and fifteen healthy women users of oral contraceptives (age: 27 ± 4 yr.; BMI: 23.0 ± 3.0 kg/m2; means ± SD) were recruited. These subjects were part of a larger study which had the following exclusion criteria: history of musculoskeletal or orthopedic injury of the spine, hip, knee, ankle, or foot; history of neurological injury or disease of the peripheral or central nervous system; current smoking habit; history of disordered eating, stress fracture, connective tissue disorder (Marfan syndrome, Ehlers-Danlos disease), or menstrual disfunction (primary or secondary amenorrhea, oligomenorrhea, anovulatory cycles, polycystic ovarian disease); current or prior pregnancy; starting or stopping OCs within the previous 6 months; use of an extended-cycle oral contraceptive that eliminates placebo pills; or use of a non-oral hormonal contraceptive. We also excluded those women exercising more than 7 hours per week or participating in competitive-level sports because of the high rate of undiagnosed menstrual dysfunction in females in this population.^12^ The Northwestern University and University of Texas Southwestern Medical Center institutional review boards approved this study.

At the time of consent, each non-user was asked to track their menses and use a home ovulation kit (Clearblue®, Procter & Gamble, USA) to estimate the length of the follicular and luteal phases and timing of ovulation. Once enrolled in the study, participants underwent a more systematic examination using every-other-day blood draws to capture the subject-specific signature of hormonal fluctuations. Non-users were scheduled to attend two testing rounds. Subjects were randomized to initiate testing in the follicular, peri-ovulatory, or luteal phase, ensuring comprehensive sampling across the menstrual cycle and enabling capture of phase transitions between consecutive cycles. Within each phase, start days varied according to participant availability, resulting in staggered sampling that increased temporal resolution across the cycle at the cohort level. One testing round was defined as the period from the start day to the same menstrual cycle day in the subsequent cycle. Hence, one testing round spanned across one menstrual cycle or two menstrual cycles depending on the starting day. Eight subjects completed only one testing round. Four subjects participated in the two testing rounds consecutively and underwent venipuncture every other day across three consecutive cycles, providing continuous longitudinal coverage across cycle boundaries.

The OC users attended one testing round and underwent venipuncture every other day for one OC pill pack. The starting date of OC users was also randomly assigned across subjects. The testing time was scheduled based on participants’ preferences and maintained at the same time of day throughout the testing sessions to minimize the effect of the diurnal fluctuation of hormone levels^13,14^. Menstrual cycle day 1 was defined as the first day of menses. Day 1 for the users was defined as the first day of active pills. The investigators were blinded to subjects’ menstrual cycle phase and contraceptive use.

Subjects were instructed to maintain stable exposure to caffeine, alcohol, and exercise for at least 12 hours before the blood draws. Venipuncture was performed on the antecubital area at each testing session using two 5 mL Vacutainer serum separator tubes (BD, Franklin Lakes, New Jersey). Once the tube was filled, the sample was mixed well and allowed to clot based on the manufacturer’s guidelines.

Systemic estradiol (E2), progesterone (P), and testosterone (T) concentrations were measured in serum. Samples obtained at Northwestern University were processed in the Shirley Ryan Ability Lab (SRA-Lab) outpatient laboratory (n non-users = 10, n users = 10). Estradiol and Progesterone levels were analyzed using the ADVIA chemiluminescence assay (Siemens ADVIA Centaur, Siemens Healthcare, Germany). Mean percentage coefficient of variation (%CV) ranged from 4-12.1% (within-run) and 4.5-8.1% (run-to-run) for Estradiol, and 2.5-12.4% (within-run) and 1.9-5.7% (run-to-run) for Progesterone. Serum testosterone was also measured using a chemiluminescent immunoassay (ADVIA Centaur, Siemens Healthcare, Germany), with reported within-run and run-to-run coefficients of variation of approximately 3–7% and 5–10%, respectively, depending on concentration range. Samples obtained at UT Southwestern Medical Center were sent to MedFusion (Lewisville, TX) on the same day of testing for processing and analysis of these hormones (n non-users = 24, n users = 5). At MedFusion, estradiol and progesterone concentrations were tested using a competitive enzyme immunoassay with no replicates. Testosterone quantification was performed using an immunochemiluminescent assay with no replicates.

Serum from the second tube was separated by centrifugation and stored at –80 °C until analysis of ethinyl estradiol (EE), progestins (SP), and relaxin-2 (RLN2). Concentrations of EE and SP were quantified using a liquid chromatography-tandem mass spectrometry method developed by our group^15^. Systemic RLN2 concentrations were measured in serum. Serum RLN2 was quantified using the MSD R-PLEX® electro-chemiluminescent assay system, which applies a highly sensitive sandwich-based detection method utilizing biotinylated capture and SULFO-TAG™ antibodies. Assays were performed according to the manufacturer’s protocol and read using the MESO QuickPlex SQ 120MM. All samples were run in duplicate, and intra- and inter-assay coefficients of variation were <10%.

### Data Analysis

The hormone profiles across the menstrual and oral contraceptive (OC) pill cycle were assessed using the R language (version 4.5.3, R Foundation for Statistical Computing, Vienna, Austria). The graphs were generated using the *plotly* package v4.12.0^16^ .

The objective of the study was to characterize subjects’ overall hormone profiles across the menstrual cycle rather than to evaluate differences between menstrual cycles. Therefore, measurements from the two rounds were combined within subject for the primary analysis. The number of participants was interpreted in the context of this high-resolution longitudinal design. The primary estimands were the shape and timing of within-cycle endocrine trajectories, rather than cross-sectional prevalence or between-group differences. Repeated every-other-day measurements across multiple cycles therefore provided dense within-person temporal information, while the hierarchical GAM explicitly accounted for dependence among repeated observations through subject-specific smooth deviations. Serial measurements were not treated as independent participants.

To characterize E2, P, and T cyclic profiles across the menstrual cycle, a generalized additive model (GAM) with subject-specific smooth effects, implemented using the *mgcv* package v1.9-4 ^17^ was fitted to model each log-transformed hormone level *Y*. GAM was selected because it enables estimation of common data patterns, which can be non-linear, without requiring a predefined parametric equation.

The model included a cyclic cubic regression spline for the normalized day (*x*) and subject-specific smooth deviations of *x* modeled using a sum-to-zero factor-smooth interaction to capture individual-level variation. To consider possible inter-platform assay bias, site of the analyses (SRA-Lab or MedFusion) was included as a fixed categorical term. The model was specified as

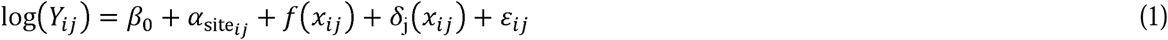

where *a*_site*ij*_ is the parametric effect of the categorical variable for analysis site (SRA-Lab or MedFusion), *f(x)* is a population-level smooth function of *x*, *o*_j_*(x)* denotes subject-specific smooth deviations. This constraint ensures that the population-level smooth *f(x)* represents the average effect across subjects, while individual curves capture deviations from this mean effect. The response distribution was considered as a Gaussian model. To characterize the population-level temporal trajectory while accounting for analysis site, predictions were generated over a dense time grid with subject-specific smooth deviations excluded. Predictions for the two levels of site were averaged equally on the linear-predictor scale. Pointwise 95% confidence intervals were calculated using the GAM coefficient covariance matrix and then exponentiated for presentation on the original response scale.

Similarly, to characterize SP, EE, and RLN2 profiles throughout the menstrual cycle, the untransformed hormone level was modeled using GAM. The model includes a cyclic cubic regression spline and subject-smooth deviations. The model can be written as

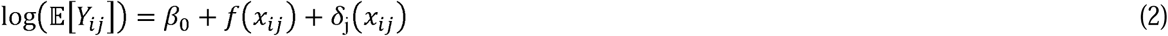

Both the global and subject-specific smooths were constrained to be cyclic over the interval [0, 1] via specified boundary knots. These analyses were conducted in-house; therefore, the categorical term for the analysis site was omitted. This model considers a Tweedie distribution with a log link because the response variable was non-negative and contained both zero and positive continuous values. The Tweedie distribution is well-suited for such data as it accommodates a compound Poisson-Gamma structure, allowing simultaneous modeling of zero observations and continuous positive responses within a unified framework. In this study, missing hormone values were not imputed. The GAM was fitted using the available outcome observations, under the assumption that missing hormone values were missing at random. For intervals where an individual had no observations, values were not imputed. Instead, the model used information from all subjects to estimate the overall time pattern, while allowing each individual to follow their own smooth trend around it, with a shared level of smoothness across individuals.

All model parameters were estimated using the restricted maximum likelihood method. Basis dimensions were chosen to be sufficiently large to satisfy model adequacy. Model adequacy was evaluated by examining residual patterns, quantile-quantile plots, and response-fitted relationships. The adequacy of the smoothing-basis dimensions was assessed using the k-index test, with values close to 1 and non-significant p-values indicating sufficient basis size. Models were considered adequate when the residuals showed no systematic patterns. Basis dimensions (*k*) for the population, smooth, and subject-specific smooths are listed in Table 2. Basis dimensions of models for non- and Table 3. Basis dimensions of GAM models for OC users..

**Table 1.** Baseline cohort characteristics.

| Characteristic | Naturally cycling women | Combined oral contraceptive users |
| --- | --- | --- |
| Participants, n | 34 | 15 |
| Age, yr, mean ± SD | 27 ± 5 | 27 ± 4 |
| BMI, kg/m<sup>2</sup>, mean ± SD | 25.4 ± 5.6 | 23.0 ± 3.0 |
| Mean menstrual-cycle length, days | 30.1 | — |

**Table 2.** Basis dimensions of models for non-users GAM.

| model | population smooth | subject-specific smooth |
| --- | --- | --- |
| E2 | 15 | 12 |
| P | 15 | 10 |
| T | 11 | 14 |
| RLN2 | 15 | 7 |

**Table 3.** Basis dimensions of GAM models for OC users.

| model | population smooth | subject-specific smooth |
| --- | --- | --- |
| E2 | 11 | 5 |
| P | 8 | 13 |
| T | 9 | 24 |
| EE | 21 | 22 |
| SP | 12 | 6 |
| RLN2 | 5 | 3 |

To determine the maximum population-level of the hormones along with their location, the fitted model, excluding subject-specific random-effect terms, was employed, yielding the marginal mean response as a function of the normalized day *x*. Population-level predictions were then obtained by applying the back-transformation to the linear predictor. Because both the population-level maximum and its corresponding location are nonlinear functionals of the fitted model, their uncertainty was quantified via Monte Carlo simulation. Coefficient vectors were sampled from a multivariate normal approximation to the posterior distribution of the model coefficients (10,000 simulations), based on the estimated covariance matrix of the fitted model, including smoothing-parameter uncertainty. Then, the population-level curve was reconstructed for each simulation, and the maximum and its location were obtained for each simulated curve. The mean and 95% confidence intervals for both the maximum value and its location were computed from the empirical distributions, using the mean and the 2.5^th^ and 97.5^th^ percentiles. For cyclic predictors, calculations were performed on an extended periodic domain to ensure correct identification of extrema near the boundary, and final estimates were mapped back to the original scale.

The location of population-level onset and offset of P and RLN2 was defined from the fitted marginal response curve obtained by excluding subject-specific terms from the model. The detection threshold was defined as the mean + 3 × standard deviation of the population-level curve at the follicular phase (normalized day 0.1 to 0.5). The onset and offset were defined as the first and last values of the normalized day *x* at which the population-level curve exceeded this threshold. Similar to the method to detect the maximum population-level, for cyclic predictors, the fitted curve was extended to ensure that threshold crossings near the boundary were correctly identified. Furthermore, the onset and offset were determined from the contiguous above-threshold segment containing the peak of the central cycle. Uncertainty in onset and offset was quantified via Monte Carlo simulation using the same approach described above (10,000 simulations), drawing coefficient vectors from a multivariate normal approximation to the posterior distribution of the model coefficients, based on the estimated covariance matrix of the fitted model, including smoothing-parameter uncertainty. To address cyclicity in the predictor, simulated onset and offset values were aligned relative to the point estimate before summarization. The mean and 95% confidence intervals were computed from the empirical distributions, using the mean and the 2.5^th^ and 97.5^th^ percentiles, and subsequently mapped back to the original cyclic domain.

## Results

Cyclic dynamics of relaxin and its temporal alignment with ovarian steroids is shown in Figure 1. To characterize endocrine coordination across the menstrual cycle, we modeled longitudinal hormone profiles using cycle-normalized time (0–1) across 80 cycles from 34 women. Testing rounds for five subjects spanned over 1 cycle, 14 subjects over 2 cycles, 13 subjects over 3 cycles, and 2 subjects over 4 cycles. Average cycle length across subjects was 30.1 days.

**Figure 1.**
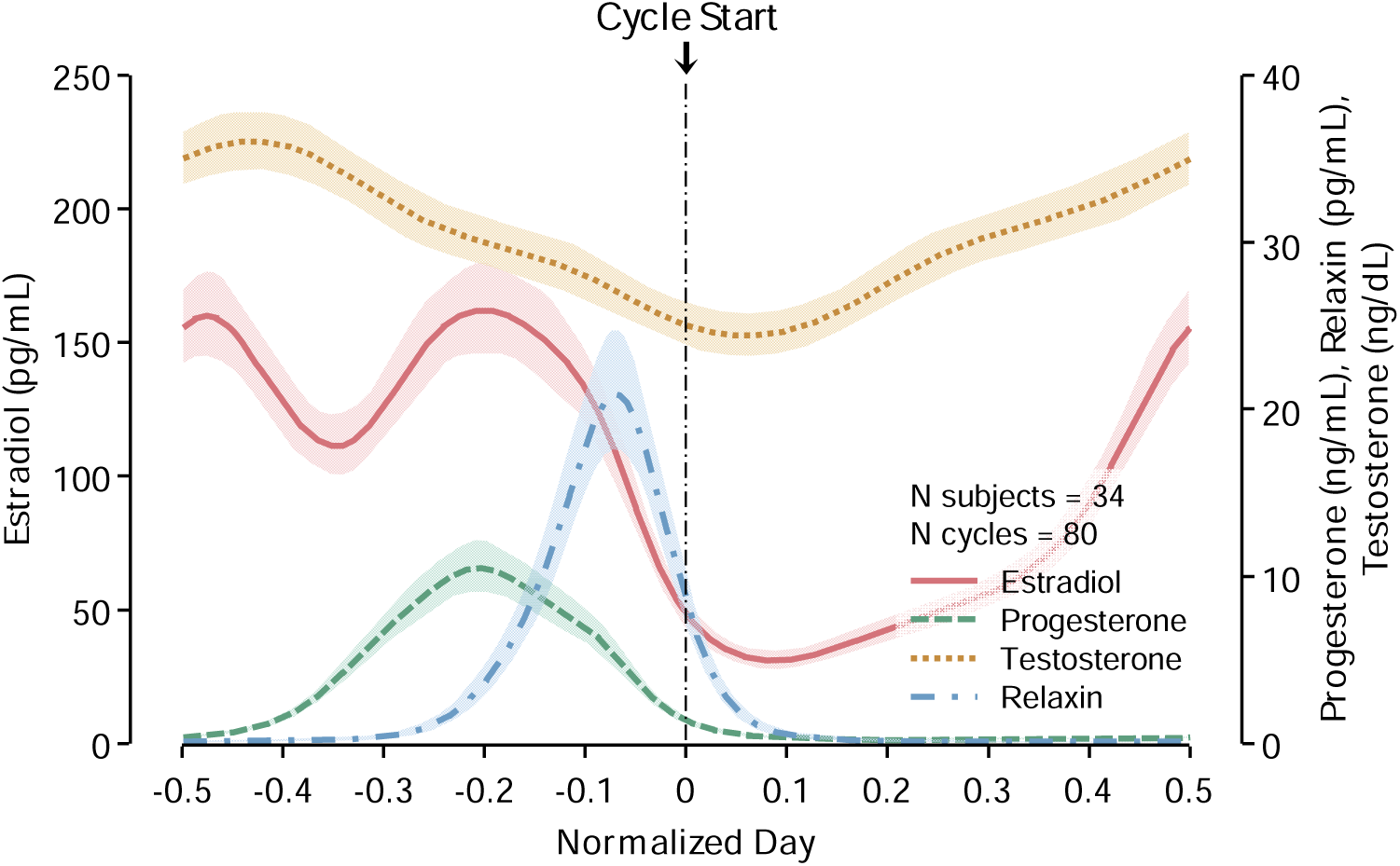
Cycle-normalized population endocrine timing architecture reconstructed from dense longitudinal measurements obtained across 80 menstrual cycles from 34 naturally cycling women. Shaded regions indicate 95% confidence intervals from hierarchical cyclic modeling. Estradiol exhibited the expected peri-ovulatory activation peak followed by a secondary luteal elevation, whereas progesterone demonstrated a broader luteal-phase activation profile. In contrast, relaxin emerged as a sharply phase-locked late-luteal endocrine transition signal with substantially lower temporal dispersion than both estradiol and progesterone. Testosterone showed comparatively modest cyclic variation. These findings identify relaxin as a previously under-resolved temporally gated system’s hormone embedded within late-luteal endocrine transition physiology.

After accounting for the overall cyclic effect of time and subject-specific deviations in the time trajectory, analysis site (SRA-Lab or MedFusion) was not significantly associated with log-transformed E2 (*/5* = 0.052, SE = 0.067, t = 0.782, p = 0.434), but statistically significantly associated with log-transformed P (*β* = -0.177, SE = 0.090, t = -1.973, p = 0.049) and T (*β* = 0.512, SE = 0.045, t = 11.26, p < 0.001).

As expected, estradiol exhibited a pre-ovulatory peak followed by a smaller luteal peak and then declined, while progesterone rose during the luteal phase. In contrast, relaxin demonstrated a distinct, sharp peak at the end of the luteal phase, temporally offset from progesterone and more narrowly distributed than estradiol and progesterone. Notably, progesterone reached its maximum shortly after the second estradiol peak in the luteal phase and before the full rise in relaxin, which might suggest that relaxin secretion is tightly coupled to corpus luteum formation events rather than ovulatory events. Testosterone exhibited relatively modest cyclic variation compared to ovarian hormones. These results establish relaxin as a phase-specific endocrine signal with high temporal precision relative to other reproductive hormones.

We next quantified the amplitude and distribution of relaxin around the beginning of the menstrual cycle (Figure 2). Relaxin concentrations were substantially higher during the late luteal phase than the first day of the menstrual cycle (mean 20.90 pg/mL [17.71–24.87] vs. 8.39 pg/mL [7.00–10.04]), corresponding to an approximate 60% relative difference when expressed as a proportion of relaxin levels. Density-based visualization revealed that relaxin exhibited a narrow distribution, whereas progesterone showed a broader profile. The temporal separation between their peaks was also evident, with relaxin peaking later and declining rapidly. These findings reinforce the concept that relaxin represents a transient but high-intensity endocrine signal distinct from the sustained luteal progesterone rise.

**Figure 2.**
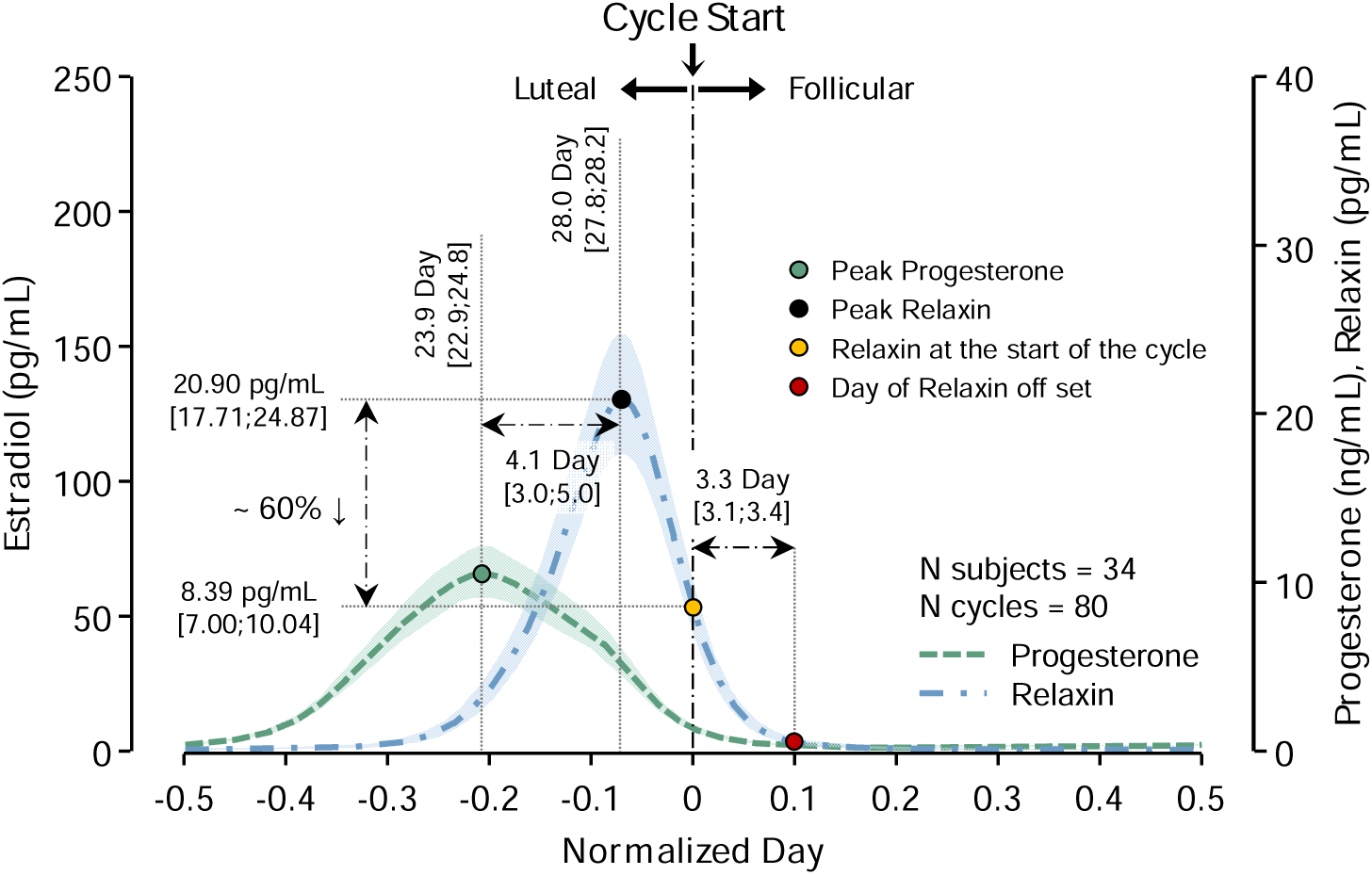
Cycle-normalized endocrine timing trajectories illustrating phase-specific endocrine dynamics and temporal dispersion across the menstrual cycle. Shaded regions represent model-estimated confidence intervals surrounding the reconstructed mean trajectories for each hormone. Vertical annotations indicate the duration and temporal positioning of major endocrine transition states, including a broader luteal activation interval (23.9 days [22.9, 24.8]) and a tightly constrained peri-transition signaling window (3.3 days [3.1, 3.4]). Colored markers denote representative endocrine states sampled along the reconstructed trajectories, highlighting progressive transition-state signaling across the cycle.

To assess the timing of peak hormone expression, we quantified the distribution of peak days across cycles (mean and 95% CI range). Progesterone peaked at day 23.9 [22.9–24.8], whereas relaxin peaked later at day 28.0 [27.8–28.2], indicating a temporal offset of ∼4.1 days between the two hormones (Figure 2). Furthermore, relative to the peak of estradiol in the follicular phase, progesterone peaked 8.2 [7.1-9.3] days later, and relaxin peaked 12.3 [11.7-13.0] days later (Figure 3). This narrow distribution of relaxin peak timing across subjects highlights its low inter-cycle variability and strong phase locking to ovulatory events and the start of the cycle, in contrast to progesterone, which demonstrated a broader distribution. These data further support the interpretation that relaxin is not simply a downstream product of progesterone-driven luteal function, but rather reflects a distinct regulatory axis linked to corpus luteum activation and late luteal transition.

**Figure 3.**
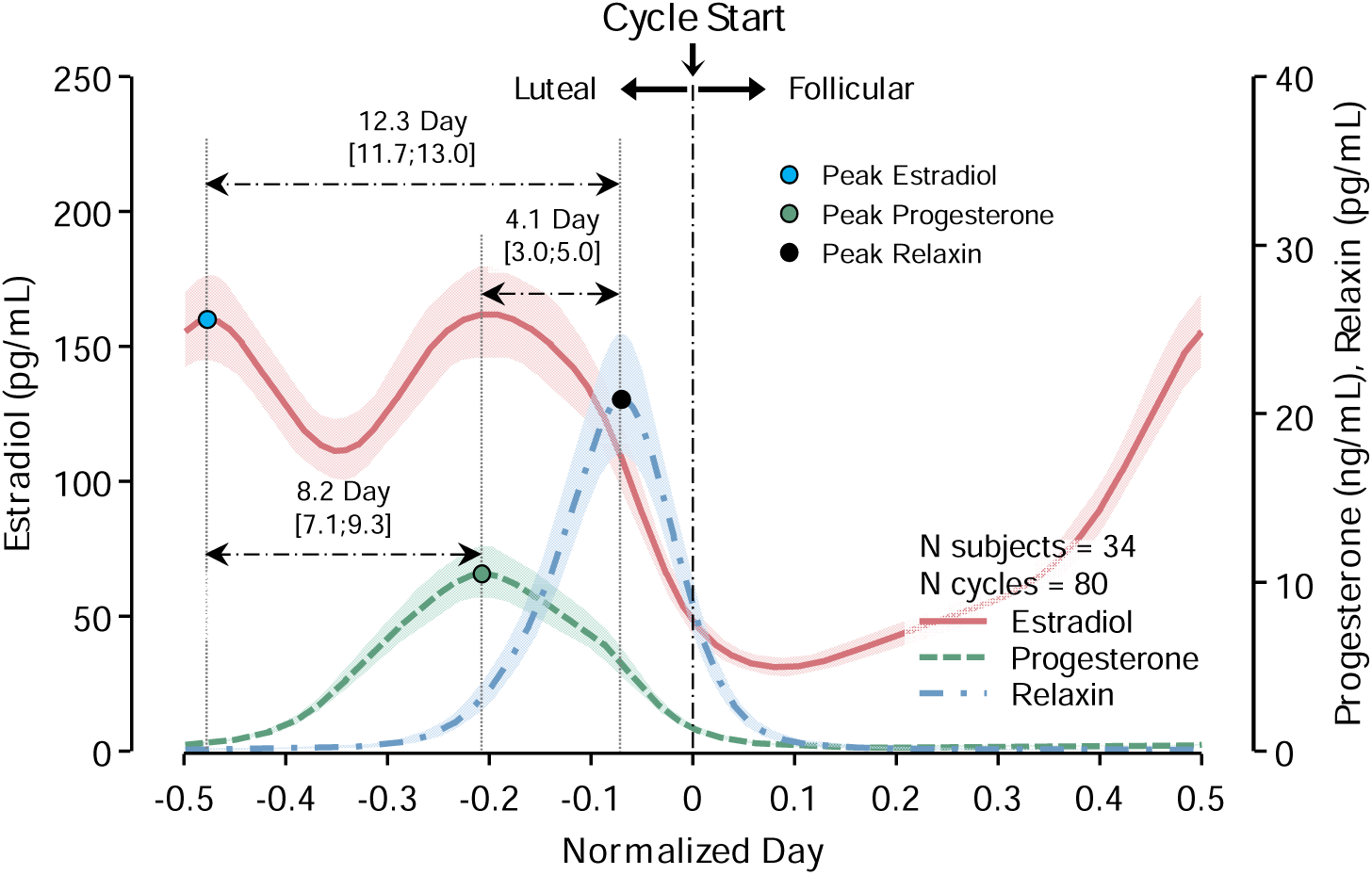
Sequential endocrine phase locking of estradiol, progesterone, and relaxin across the menstrual cycle. Shaded regions indicate 95% confidence intervals from hierarchical cyclic modeling. Relative to the follicular estradiol peak, progesterone peaked 8.2 days later, whereas relaxin peaked 12.3 days later. Relaxin, therefore, followed both estradiol and progesterone within a highly reproducible endocrine sequence consistent with delayed corpus luteum–dependent transition-state signaling. These findings support a temporally staged endocrine control architecture rather than simultaneous luteal hormone release.

The cohort-level model derived from 34 subjects across 80 cycles defines the population-average endocrine trajectory across normalized menstrual time, leveraging randomized phase initiation and staggered entry days to achieve dense temporal coverage at the group level (Figure 1). In contrast, the subset of four women followed every other day across two consecutive cycles beginning at the early luteal phase preserves within-subject continuity across cycle boundaries and is therefore particularly informative for assessing cycle-to-cycle reproducibility and transitional dynamics at menstruation (Figure 4). Accordingly, the larger cohort model is best interpreted as the canonical cycle-normalized waveform, whereas the back-to-back subset provides insight into recurrence, reset behavior, and possible short-lived persistence of hormone signals across adjacent cycles. Together, these analyses strengthen inference by linking population-level temporal structure to directly observed within-subject cycle continuity.

**Figure 4.**
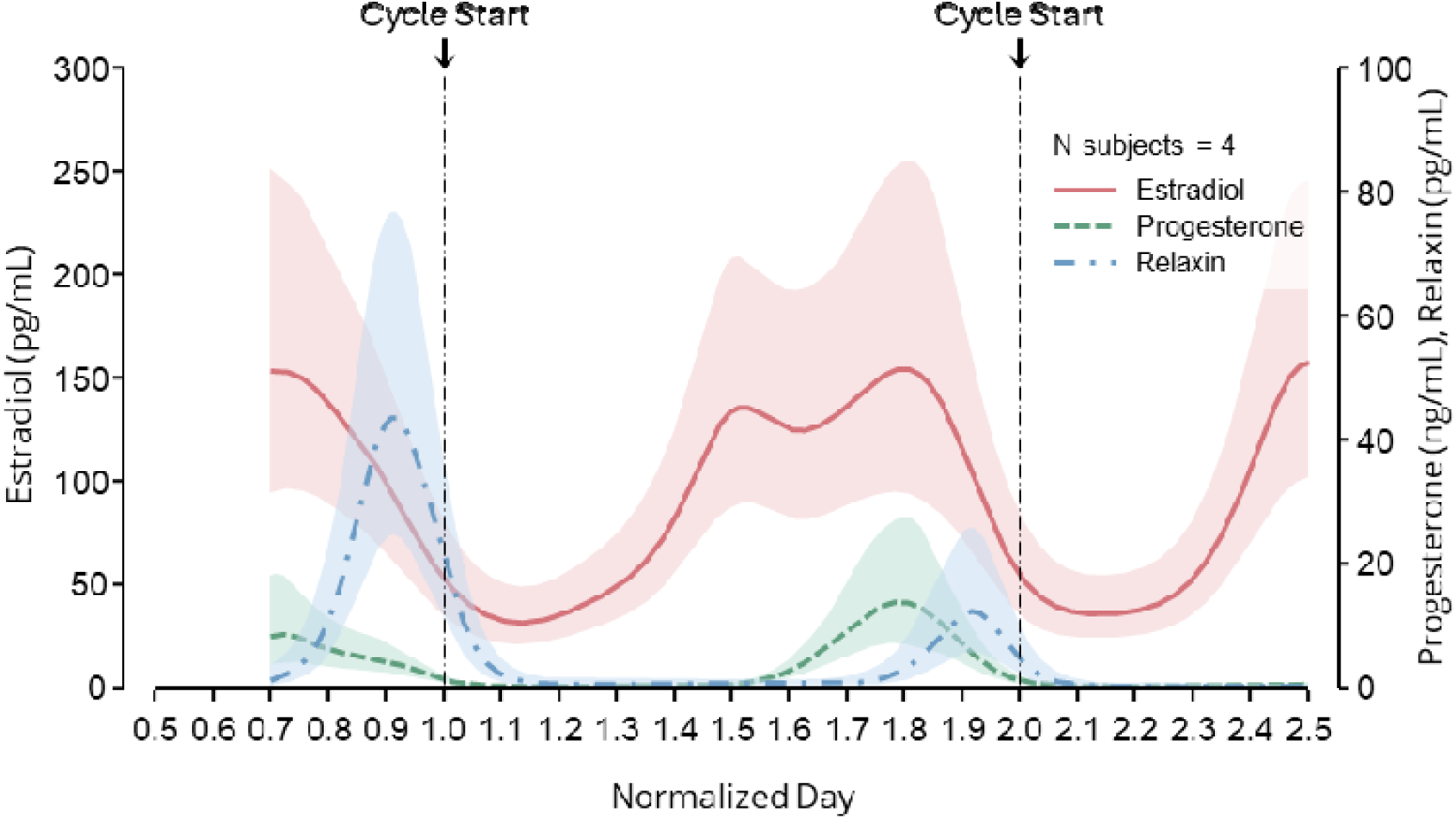
Consecutive-cycle longitudinal endocrine trajectories from four women sampled every other day across two sequential menstrual cycles beginning in the luteal phase. Shaded regions indicate 95% confidence intervals from hierarchical cyclic modeling. Repeated phase-locked late-luteal relaxin activation and subsequent decline across the menstrual boundary were preserved within individuals, supporting strong cycle-to-cycle reproducibility of the population-derived endocrine timing architecture. These data complement cohort-level modeling by directly demonstrating continuity and recurrence of temporally gated relaxin signaling within subjects.

To evaluate how sampling structure influences observed hormone distributions, we compared model-derived trajectories with raw, non-normalized data from 80 cycles. Violin plots of observed values demonstrated substantially greater dispersion, particularly for relaxin, consistent with phase misalignment and sparse sampling across subjects (Figure 5). Despite this increased variability, the relative ordering of hormone magnitudes was preserved, with relaxin maintaining a distinct distribution from progesterone.

**Figure 5.**
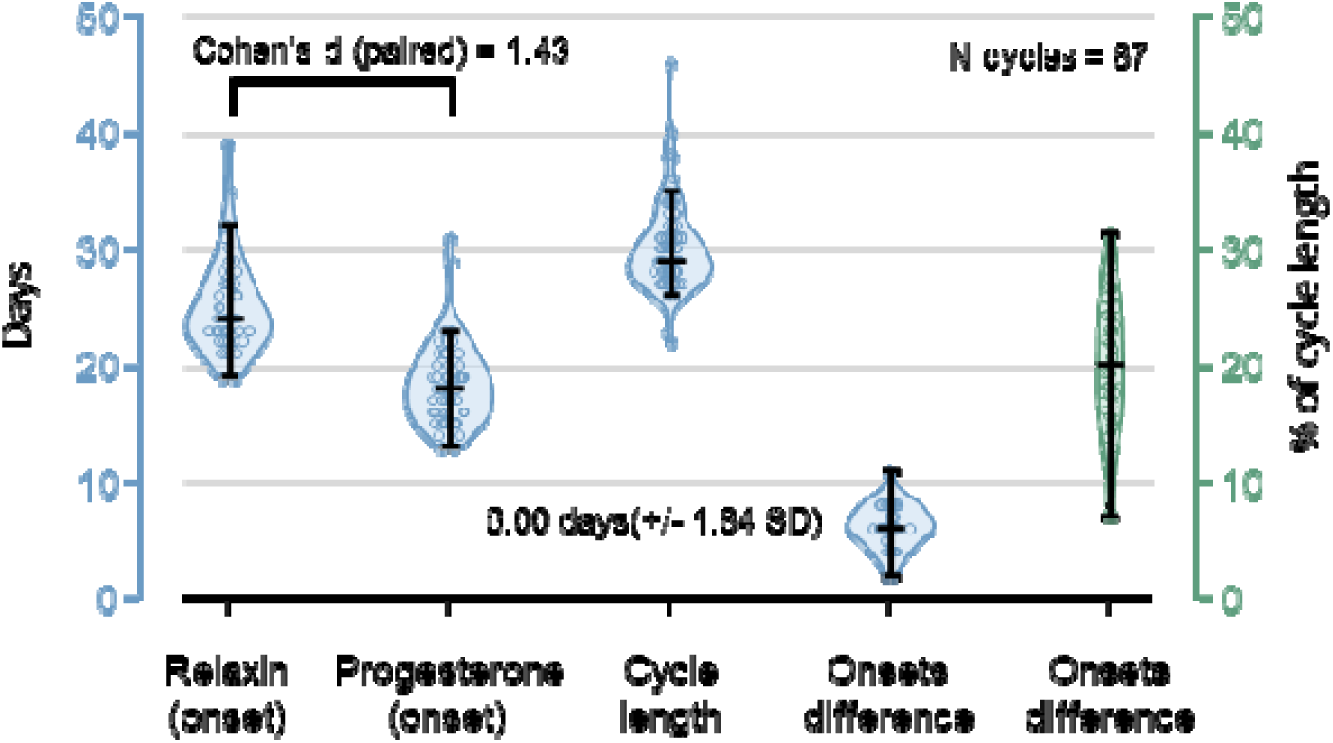
Distributional analysis of endocrine onset timing across 80 menstrual cycles. Relaxin onset occurred 6.0 ± 1.84 days later than progesterone onset, with a large, paired effect size (Cohen’s d = 1.43). This reproducible temporal delay indicates that relaxin is not synchronously released with progesterone despite their shared luteal origin, supporting a temporally staged corpus luteum endocrine control architecture with distinct transition-state signaling phases.

For the non-users, two complementary representations of the data are presented. The model-based analysis (Figure 1) uses cycle normalization and continuous function estimation to align subjects across a standardized temporal axis (0–1), thereby recovering the underlying physiological trajectory of the variable across the menstrual cycle. In contrast, the raw data (Figure 5) reflects discrete measurements obtained every other day without phase alignment, with variable cycle start points across subjects. As a result, the raw distributions incorporate phase misalignment, sparse sampling, and inter-individual variability, which collectively broaden distributions and attenuate apparent peak differences. The model, therefore, provides an estimate of the latent cycle-dependent signal, whereas the raw data represents sampled observations of that signal under real-world constraints. The differences between the two figures are thus expected and reflect methodological rather than biological discrepancies.

In women using combined oral contraceptives (OCPs), cyclic variation in endogenous hormones was markedly attenuated (Figure 6). Ethinyl estradiol and progestin levels remained relatively stable across the cycle, and importantly, relaxin production was effectively absent. This suppression of relaxin is consistent with inhibition of ovulation and corpus luteum formation under OCP use, supporting the interpretation that relaxin production is dependent on ovulatory and luteal processes rather than circulating progestin exposure. These findings further argue against a simple progesterone-driven model of relaxin regulation and instead support a source-dependent mechanism linked to corpus luteum function.

**Figure 6.**
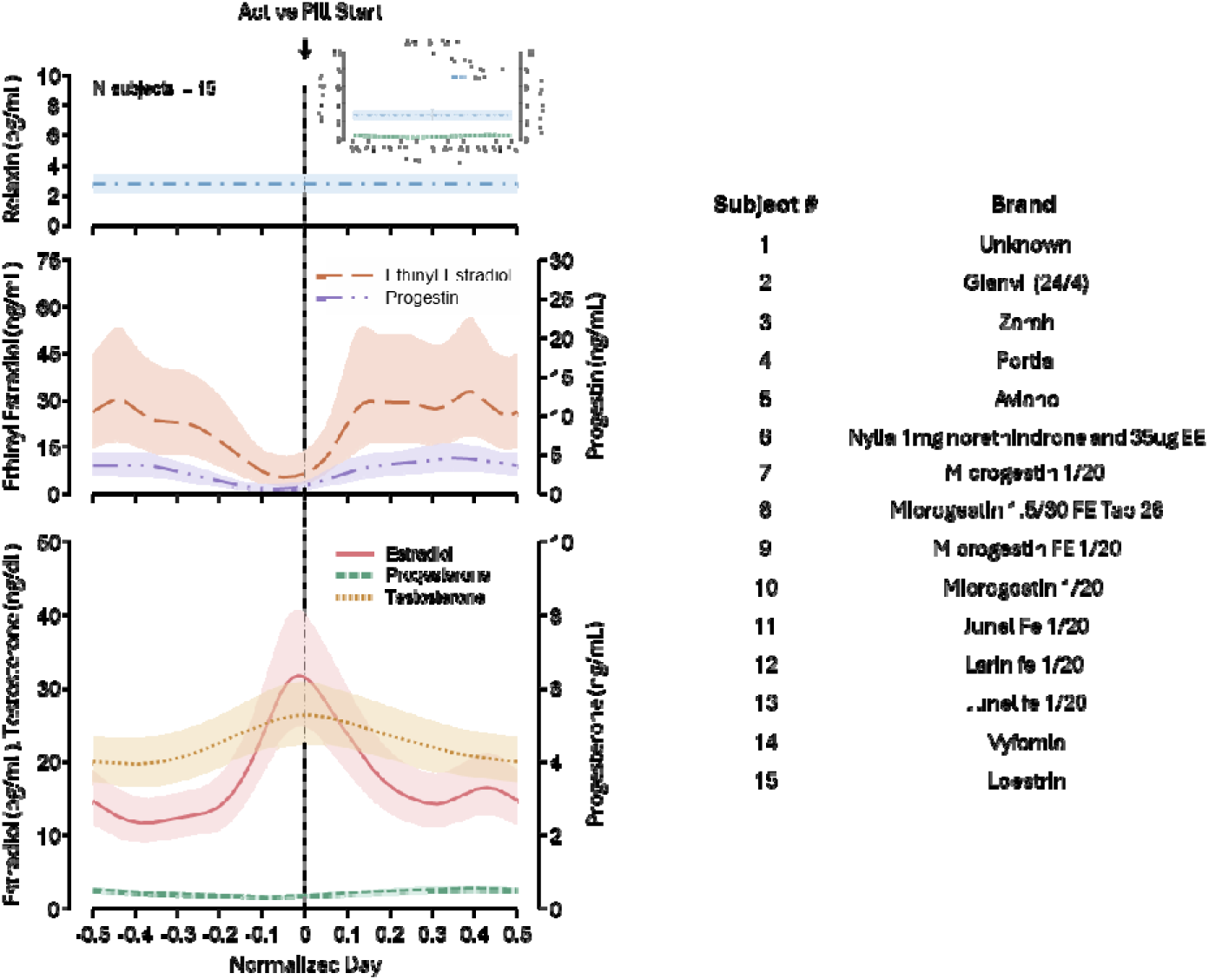
Cycle-normalized endocrine timing architecture from 15 women using combined oral contraceptives, along with a table listing the commercial brand names of contraceptives used b participants. Endogenous estradiol and progesterone cyclicity was markedly attenuated, whereas ethinyl estradiol and progestin concentrations reflected externally imposed pill-phase exposure. Circulating relaxin was nearly absent throughout the cycle despite sustained exogenous progestin exposure. Thes findings indicate that relaxin signaling depends on ovulation-linked corpus luteum transition biology rather than steroid exposure alone, further supporting relaxin as a structurally gated endocrine signal.

## Discussion

In this study, we demonstrate that relaxin exhibits a consistent luteal-phase rise in nonpregnant women, occurring in close temporal association with progesterone but with a characteristic delay in both onset and peak. These findings confirm that relaxin is an active luteal hormone in the natural menstrual cycle and refine its temporal profile with greater precision than previously reported.

### Dense longitudinal design

An important strength of the present study is the temporal density of sampling across the late-luteal-to-menstrual transition. Although the number of unique participants is modest, the primary objective was to characterize the shape, onset, peak, offset, and relative timing of within-cycle endocrine trajectories rather than population prevalence or cross-sectional differences. Participants underwent approximately every-other-day venipuncture across 80 menstrual cycles. For comparison, the U.S. BioCycle Study ^18^ followed 259 healthy premenopausal women using up to eight hormonally targeted visits per cycle. Visit timing was initially individualized according to each participant’s self-reported usual cycle length and subsequently adjusted around the periovulatory interval using urinary fertility-monitor measurements. Importantly, BioCycle reported that self-reported cycle length differed from the subsequently observed cycle length by three or more days in approximately 40% of cycles, and cycle length differed by three or more days between consecutive cycles in nearly half of women. Thus, although fertility monitoring provided an effective prospective anchor for periovulatory sampling, the timing of the subsequent menstrual boundary could not be known prospectively. This distinction is particularly relevant for relaxin, whose narrow rise and peak occur near the late-luteal-to-menstrual transition. The every-other-day design used here provides multiple observations across this interval without requiring prospective prediction of the eventual cycle length, allowing endocrine trajectories to be aligned subsequently to the observed menstrual boundary. The hierarchical modeling framework further accounts for the repeated-measures structure through subject-specific smooth effects, such that serial observations increase information regarding within-person temporal trajectories without being treated as independent participants.

An additional attribute of the present design is that participants did not enter the study at a uniform or prespecified menstrual-cycle day. Instead, sampling was initiated at staggered points across the follicular, peri-ovulatory, and luteal phases and then continued approximately every other day – at random across subjects based on the participants’ availability. This reduced dependence on prospective assumptions regarding cycle length or the precise timing of endocrine events and provided distributed temporal coverage across the menstrual cycle at the cohort level. Moreover, several participants were followed continuously across consecutive cycles, allowing direct observation of cycle-to-cycle recurrence and endocrine behavior across the menstrual boundary. This design is particularly relevant for relaxin, whose narrow late-luteal rise and persistence into early menstruation may be difficult to resolve using sparse phase-targeted sampling. Together, the staggered-entry and consecutive-cycle strategy provided both broad cohort-level temporal coverage and within-person information regarding the reproducibility of the observed endocrine trajectory.

### Progesterone–relaxin timing and its physiological relevance

In humans and other primates, the corpus luteum is a key source of relaxin. The control architecture appears, however, more complex and less progesterone-dependent than a simple source-driven model would predict^1^. While plausible, primate *in vivo* studies demonstrate a dissociation between progesterone depletion and relaxin secretion, indicating that progesterone is unlikely to serve as the sole or dominant proximal driver of relaxin output ^19,20^. At the gene-regulatory level, progesterone can influence relaxin biology. A human cell model study reported that a progesterone/glucocorticoid-responsive element in the relaxin gene regulatory regions was functional. However, significant, this cell-level finding does not prove *in vivo* human luteal control, but it does support the narrower idea that progesterone-family signaling can regulate relaxin gene expression under some cellular contexts^21^. Collectively, these observations argue that progesterone contributes to the broader luteal regulatory milieu but does not serve as the primary proximal driver of relaxin production, which instead appears to be governed by luteal structure, differentiation state, and gonadotropic signaling.

The absence of circulating relaxin in women taking combined oral contraceptives (Figure 5), despite exposure to exogenous progestins, it provides important insight into the regulation of relaxin production. Oral contraceptives suppress the hypothalamic–pituitary–ovarian axis and prevent the LH surge required for ovulation, thereby eliminating corpus luteum formation and the associated luteal endocrine environment. As a result, hormone production is normally dependent on luteal structure, including progesterone and other luteal-derived factors, which are fundamentally altered or absent, despite the presence of exogenous steroid hormones. The absence, however, of relaxin in women on oral contraceptives is not evidence against hormonal regulation per se but rather reflects the absence of the corpus luteum—the primary structural and endocrine source required for relaxin production.

The observed lag of approximately 6 days between the rise of progesterone and relaxin, and the ∼4-day separation between their peaks, aligns with classic endocrine studies showing that progesterone increases immediately post-ovulation, whereas relaxin secretion occurs later in the luteal phase ^3,7,8^. This temporal offset likely reflects normal luteal maturation: progesterone initiates secretory differentiation and immune modulation of the endometrium, while relaxin, produced by the same corpus luteum, supports subsequent vascular, stromal, and extracellular-matrix remodeling required for adequate luteal function ^6,22,23^.

Vascular function varies significantly across the menstrual cycle, with human studies demonstrating cyclic changes in endothelial function, arterial compliance, and hemodynamics ^24–26^. Flow-mediated dilation and endothelial reactivity increase during the late follicular phase, decline in the early luteal phase, and partially recover in the late luteal phase ^25,27^. These physiological changes are accompanied by measurable fluctuations in systemic blood pressure and nitric oxide bioavailability, with even modest cycle-dependent reductions in blood pressure associated with meaningful reductions in cardiovascular risk ^28,29^. While estradiol-mediated nitric oxide signaling explains early-cycle vascular improvements ^30,31^, the persistence and recovery of vascular function in the luteal phase are not fully accounted for by estradiol alone ^25,26^. Relaxin, which peaks in the late luteal phase and promotes nitric oxide production, vascular remodeling, and arterial compliance, provides a plausible mechanistic link between endocrine state and vascular structure ^32–34^. Integrating relaxin into models of menstrual cycle physiology thus offers a more complete, state-dependent framework for understanding cardiovascular dynamics in premenopausal women.

The delayed relaxin rise also has implications for connective-tissue physiology. Musculoskeletal performance and injury risk in women exhibit clear cycle-dependent variation, with changes in strength, joint laxity, and ligament integrity reported across the menstrual cycle ^35^. While these effects are often attributed to estradiol and progesterone, emerging evidence suggests that vascular and extracellular matrix dynamics also play a critical role. Endothelial function and blood flow vary across the cycle, influencing tissue perfusion and fatigue resistance, while hormone-dependent remodeling of connective tissue alters stiffness and force transmission. Recent high-level evidence synthesis concluded that, among the female sex hormones studied, 17β-estradiol and relaxin appear to exert the greatest effects on both the mechanical and cellular properties of ligaments, supporting a plausible endocrine basis for connective tissue variability across reproductive states ^36^. However, inter-individual responsiveness remains poorly defined. These observations suggest that Relaxin is a biologically plausible and understudied mediator of hormone-state-dependent connective tissue. By modulating both vascular delivery and connective tissue properties, relaxin may contribute to cycle-dependent variability in performance and injury susceptibility, offering a unified framework linking cardiovascular physiology to neuromechanical output. By defining when relaxin peaks relative to progesterone, our findings help clarify the timing of this vulnerability window within the menstrual cycle.

### Physiological and clinical implications of post-luteal relaxin persistence

Measurable relaxin concentrations were observed in the early menstrual phase, consistent with the gradual luteal regression and continued release of stored peptide despite declining steroid secretion ^1,3^. Although transient, this “relaxin tail” may influence several physiological processes. Relaxin modulates matrix turnover by upregulating matrix metalloproteinases (MMPs) and reducing collagen and TIMP expression in endometrial stromal cells ^22,23,37^. Such actions may facilitate coordinated tissue breakdown and early re-epithelialization during menstruation. Insufficient relaxin exposure could theoretically impair this repair process or contribute to abnormal uterine bleeding, although formal clinical evidence is limited.

Relaxin also enhances vasodilatory pathways, notably via NO- and VEGF-dependent mechanisms^2,6^. Relaxin persistence into menses may therefore modulate uterine perfusion and menstrual discomfort. Dysmenorrhoea is associated with prostaglandin-mediated vasoconstriction and uterine ischemia^38^. Reduced relaxin activity during this transition may exacerbate ischemic pain, whereas prolonged relaxin exposure may offer a modest protective effect. Additionally, because relaxin affects stromal compliance and angiogenesis in ovarian and uterine tissues, its presence across the menstrual boundary may contribute to cycle-to-cycle physiological continuity, with possible subtle effects on early follicular microvascular dynamics. These hypotheses remain speculative but are consistent with known relaxin biology.

Together, these findings refine the temporal architecture of relaxin secretion during the luteal phase and demonstrate a reproducible temporal relationship between relaxin and progesterone. The persistence of measurable relaxin into early menstruation further identifies a previously under-resolved endocrine transition across the luteal–menstrual boundary. Although relaxin has established actions in vascular and connective-tissue biology, the present study did not directly assess endometrial function, tissue perfusion, menstrual symptoms, or musculoskeletal outcomes. Accordingly, the physiological consequences of this late-luteal and early menstrual relaxin signal remain to be determined. These data therefore provide a temporal framework for future studies testing whether variation in relaxin timing or magnitude is associated with reproductive, vascular, or musculoskeletal physiology across the menstrual cycle.

### Limitations

This study provides, to our knowledge, the first cycle-resolved quantification of relaxin and progesterone profiles in nonpregnant women using dense, multi-day sampling across multiple cycles. While this design offers unprecedented temporal resolution, several limitations warrant consideration.

First, although the sampling frequency was sufficient to define the relative timing of progesterone and relaxin, continuous daily sampling was logistically complex. As a result, the precise onset and peak values, while modeled robustly, may still reflect interpolation across missing days. Future studies using fully daily sampling or automated hormonal monitoring technologies would further refine the temporal precision demonstrated here.

Second, the study relied on circulating hormone concentrations, which reflect integrated endocrine output but cannot directly identify tissue-specific sources or actions. Complementary approaches—including measurement of luteal blood flow, corpus luteum vascularization, corpus luteum transcriptomics, or endometrial tissue biomarkers—would help clarify the mechanisms underlying the progesterone–relaxin temporal offset and its inter-individual variability.

Third, the number of unique participants should be interpreted in the context of the study’s dense longitudinal design. The primary objective was estimation of within-cycle endocrine trajectories and relative hormone timing rather than population prevalence or subgroup effects. Every-other-day sampling across 80 menstrual cycles provided substantially greater temporal information per participant than conventional phase-based sampling and supported estimation of both population-level and subject-specific trajectories. Nevertheless, the number of independent participants limits the ability to evaluate demographic and clinical moderators of relaxin secretion, including BMI, parity, physical activity, and subtle luteal-phase abnormalities. Larger cohorts will therefore be required specifically for between-person and multivariable analyses.

Fourth, functional correlates such as endometrial perfusion, menstrual symptoms, or musculoskeletal outcomes were not assessed in parallel with hormone profiles. Integrating endocrine profiling with physiological measurements—including uterine blood flow, ligament compliance, or pain indices—would help test the mechanistic hypotheses generated by our findings.

Finally, assays for peptide hormones such as relaxin remain technically challenging. Although quality controls were rigorous, assay variability and low-level measurements near the detection limit may influence the characterization of early-menses relaxin persistence. Future analyses incorporating ultra-sensitive immunoassays or mass-spectrometric approaches could further strengthen confidence in low-concentration measurements.

Despite these limitations, this study provides the most detailed characterization to date of relaxin secretion across natural menstrual cycles and establishes a quantitative framework for future mechanistic and translational investigations.

## Conclusion

Using dense longitudinal endocrine phenotyping across 80 menstrual cycles from healthy women, combined with probabilistic nonlinear dynamical modeling, we identify a previously unresolved temporal architecture of relaxin signaling during the human menstrual cycle. We show that relaxin exhibits a sharply constrained late-luteal surge with substantially lower temporal dispersion than classical ovarian steroids, peaks approximately four days after progesterone, and persists into early menstruation as a post-luteal “relaxin tail” that bridges endocrine states across consecutive cycles. In women using combined oral contraceptives, endogenous relaxin was nearly absent despite sustained exogenous progestin exposure, indicating that relaxin secretion depends on ovulation-linked corpus luteum biology rather than steroid exposure alone.

Collectively, these findings redefine relaxin from a hormone primarily associated with pregnancy to a dynamic regulator of normal cyclic female physiology. The work further suggests that the menstrual cycle contains temporally staged endocrine transition states rather than static hormonal phases, with relaxin functioning as a delayed systems-level mediator potentially involved in vascular adaptation, extracellular matrix remodeling, inflammatory regulation, and menstrual repair.

More broadly, the study demonstrates how dense longitudinal phenotyping and probabilistic temporal modeling can reveal previously hidden endocrine dynamics that remain unresolved using conventional sparse sampling approaches. We believe the manuscript will be of broad interest to readers in endocrinology, systems physiology, women’s health, reproductive biology, vascular biology, and computational medicine.

## CODE AVAILABILITY STATEMENT

All analyses were conducted in R (version 4.5.3) using the mgcv library for hierarchical generalized additive modeling. Custom scripts for cycle normalization, Monte Carlo uncertainty propagation, and trajectory reconstruction are available from the corresponding author upon reasonable request and will be placed on GitHub/Zenodo.

## DATA AVAILABILITY STATEMENT

The data supporting the findings of this study are available from the corresponding author upon reasonable request. Processed longitudinal endocrine trajectories, summary statistics, and model-derived outputs necessary to reproduce the principal findings are available within the manuscript and Supplementary Information. Source data underlying the figures will be provided upon publication.

## AUTHOR CONTRIBUTIONS

### Author Contribution

Y.D. Conceived study, designed analyses, interpreted data, wrote manuscript

S.S. Statistical modeling, data processing, computational analysis

S.L. Sample preparation and methodological support

L.A.R. Hormone assay expertise and oral contraceptive endocrine analysis

E.C. Clinical interpretation and study oversight

All authors reviewed and approved the final manuscript.

